# LOTUS: a Single- and Multitask Machine Learning Algorithm for the Prediction of Cancer Driver Genes

**DOI:** 10.1101/398537

**Authors:** Olivier Collier, Véronique Stoven, Jean-Philippe Vert

## Abstract

Cancer driver genes, i.e., oncogenes and tumor suppressor genes, are involved in the acquisition of important functions in tumors, providing a selective growth advantage, allowing uncontrolled proliferation and avoiding apoptosis. It is therefore important to identify these driver genes, both for the fundamental understanding of cancer and to help finding new therapeutic targets. Although the most frequently mutated driver genes have been identified, it is believed that many more remain to be discovered, particularly for driver genes specific to some cancer types.

In this paper we propose a new computational method called LOTUS to predict new driver genes. LOTUS is a machine-learning based approach which allows to integrate various types of data in a versatile manner, including informations about gene mutations and protein-protein interactions. In addition, LOTUS can predict cancer driver genes in a pan-cancer setting as well as for specific cancer types, using a multitask learning strategy to share information across cancer types.

We empirically show that LOTUS outperforms three other state-of-the-art driver gene prediction methods, both in terms of intrinsic consistency and prediction accuracy, and provide predictions of new cancer genes across many cancer types.

**Author summary:** Cancer development is driven by mutations and dysfunction of important, so-called cancer driver genes, that could be targeted by targeted therapies. While a number of such cancer genes have already been identified, it is believed that many more remain to be discovered. To help prioritize experimental investigations of candidate genes, several computational methods have been proposed to rank promising candidates based on their mutations in large cohorts of cancer cases, or on their interactions with known driver genes in biological networks. We propose LOTUS, a new computational approach to identify genes with high oncogenic potential. LOTUS implements a machine learning approach to learn an oncogenic potential score from known driver genes, and brings two novelties compared to existing methods. First, it allows to easily combine heterogeneous informations into the scoring function, which we illustrate by learning a scoring function from both known mutations in large cancer cohorts and interactions in biological networks. Second, using a multitask learning strategy, it can predict different driver genes for different cancer types, while sharing information between them to improve the prediction for every type. We provide experimental results showing that LOTUS significantly outperforms several state-of-the-art cancer gene prediction softwares.

## Introduction

In our current understanding of cancer, tumors appear when some cells acquire functionalities that give them a selective growth advantage, allowing uncontrolled proliferation and avoiding apoptosis [1, 2]. These malignant characteristics arise from various genomic alterations including point mutations, gene copy number variants (CNVs), translocations, inversions, deletions, or aberrant gene fusions. Many studies have shown that these alterations are not uniformly distributed across the genome [3, 4], and target specific genes associated with a limited number of important cellular functions such as genome maintenance, cell survival, and cell fate [5]. Among these so-called *driver genes*, two classes have been distinguished in the literature: *tumor suppressors genes* (TSGs) and *oncogenes* (OGs) [6, Chapter 15]. TSGs, such as TP53 [7], participate in defense mechanisms against cancer and their inactivation by a genomic alteration can increase the selective growth advantage of the cell. On the contrary, alterations affecting OGs, such as KRAS [8] or ERBB2 [9], can be responsible for the acquisition of new properties that provide some selective growth advantage or the ability to spread to remote organs. Identifying driver genes is important not only from a basic biology point of view to decipher cancer mechanisms, but also to identify new therapeutic strategies and develop precision medicine approaches targeting specifically mutated driver genes. For example, Trastuzumab [10] is a drug given against breast cancer that targets the protein precisely encoded by ERBB2, which has dramatically improved the prognosis of patients whose tumors overexpress that OG.

Decades of research in cancer genomics have allowed to identify several hundreds of such cancer genes. Regularly updated databases such as the Cancer Gene Census (CGC) [11], provide catalogues of genes likely to be causally implicated in cancer, with various levels of experimental validations. Many cancer genes have been identified recently by systematic analysis of somatic mutations in cancer genomes, as provided by large-scale collaborative efforts to sequence tumors such as The Cancer Genome Atlas (TCGA) [12] or the International Cancer Genome Consortium (ICGC) [13]. Indeed, cancer genes tend to be more mutated than non-cancer genes, providing a simple guiding principle to identify them. In particular, the COSMIC database [14] is the world’s largest and most comprehensive resource of somatic mutations in coding regions. It is now likely that the most frequently mutated genes have been identified [15]. However, the total number of driver genes is still a debate, and many driver genes less frequently mutated, with low penetrance, or specific to a given type of cancer are still to be discovered.

The first methods to identify driver genes from catalogues of somatic mutations simply compared genes based on somatic mutation frequencies, which was proved to be far too basic [16]. Indeed, mutations do not appear uniformly on the genome: some regions of the genome may be more affected by errors because they are more often transcribed, so that some studies actually overestimated the number of driver genes because they were expecting lower mutation rates than in reality. Mathematically, they were formulating driver prediction as a hypothesis testing problem with an inadequate null hypothesis [17]. Several attempts have been made to adequately calibrate the null hypothesis, like [16] or [18], where it is assumed that mutations result from a mixture of several mutational processes related to different causes.

A variety of bioinformatics methods have then been developed to complete the list of pan-cancer or cancer specific driver genes. Globally, they fall into three main categories. First, a variety of “Mutation Frequency” methods such as MuSiC [19] or ActiveDriver [20] identify driver genes based on the assumption that they display mutation frequencies higher than those of a background mutation model expected for passenger mutations. However, this background rate may differ between cell types, genome positions or patients. In order to avoid such potential bias, some methods like MutSigCV [21] derive a patient-specific background mutation model, and may take into account various criteria such as cancer type, position in the genome, or clinical data. Second, “Functional impact” methods such as OncodriveFM [22] assume that driver genes have higher frequency of mutations expected to impact the protein function (usually missense mutations) than that observed in passenger genes. Third, “Pathway-based” methods consider cancer as a disease in which mutated genes occupy key roles in cancer-related biological pathways, leading to critical functional perturbations at the level of networks. For example, DriverNet [23] identifies driver genes based on their effect in the transcription networks. Although these methods tend to successfully identify the most frequently mutated genes, their overall prediction overlap is modest. Since they rely on complementary statistical strategies, one could recommend to use them in combination. The results of some of these tools are available at the Driver DB database [24].

Some methods integrate information on mutation frequency and functional impact of mutations, or other types of data such as genome position, copy number variations (CNVs) or gene expression. The underlying idea is that combining data should improve the prediction performance over tools that use a single type of information. For example, TUSON [25] or DOTS-Finder [26] combine mutation frequencies and functional impact of mutations to identify OGs and TSGs. Also in this category, the 20/20+ method [27] encodes genes with features based on their frequency and mutation types, in addition to other biological information such as gene expression level in difference cancer cell lines [28] or replication time. Then, 20/20+ predicts driver genes with a random forest algorithm, which constitutes the first attempt to use a machine learning method in this field. In [27], the authors benchmark 8 driver gene prediction methods based on several criteria including the fraction of predicted genes in CGC, the number of predicted driver genes and the consistency. Three methods proved to perform similarly on all criteria, and better than the five others: TUSON, MutSigCV, and 20/20+, validating the relevance of combining heterogeneous information to predict cancer genes.

In the present paper, we propose a new method for cancer driver gene prediction called *Learning Oncogenes and TUmor Suppressors* (LOTUS). Like 20/20+, LOTUS is a machine learning-based method, meaning that it starts from a list of known driver genes in order to “learn” the specificities of such genes and to identify new ones. In addition, LOTUS presents two unique characteristics with respect to previous work in this field. First, it combines informations from all three types of informations likely to contain information to predict cancer genes (mutation frequency, functional impact, and pathway-based informations). This integration of heterogeneous informations is carried out in a unified mathematical and computational framework thanks to the use of kernel methods [29], and allows in principle to integrate other sources of data if available, such as transcriptomic or epigenomic information. More precisely, in our implementation we predict cancer driver genes based not only on gene mutations features like “Mutation Frequency” and “Functional Impact” methods do, but also on known protein-protein interaction (PPI) network like “Pathway-based” methods do. Indeed, the use of PPI information is particularly relevant since it has been reported that proteins encoded by driver genes are more likely to be involved in protein complexes and share higher “betweenness” than a typical protein [25]. Second, LOTUS can predict cancer genes in a pan-cancer setting, as well as for specific cancer types, using a multitask learning strategy [30]. The pan-cancer setting has been adopted by most available prediction methods, since more data is available when pooling together all cancer types. The cancer type-specific prediction problem has been less explored so far, because the number of known driver genes for a given cancer is often too small to build a reliable prediction model, and because the amount of data such as somatic mutations to train the model is smaller than in the pan-cancer setting. However, the search for cancer specific driver genes is relevant, because cancer is a very heterogeneous disease: different tumorigenic processes seem to be at work in different tissue types, and consequently every cancer type probably has its own list of driver genes [15]. LOTUS implements a multitask algorithm that predicts new driver genes for a given cancer type based on its known driver genes, while also taking into account the driver genes known for other types of cancers according to their similarities with the considered type of cancer. Such approaches are of particular interest when the learning data are scarce in each individual tasks: they increase the amount of data available for each task and thus perform statistically better. To our knowledge, while a similar approach was used to predict disease genes across general human diseases [31], this is the first time a multitask machine learning algorithm is used for the prediction of cancer driver genes.

We compare LOTUS to the three best state-of-the art cancer prediction methods according to [27]. We show that that LOTUS outperforms the state-of-the-art in its ability to identify novel cancer genes, and clarify the benefits of heterogeneous data integration and of the multitask learning strategy to predict cancer type-specific driver genes. Finally, we provide predictions of new cancer genes according to LOTUS, as well as supporting evidence that those predictions are likely to contain new cancer genes.

## Results

### LOTUS, a new method for pan-cancer and cancer specific driver gene prediction

We propose LOTUS, a new method to predict cancer driver genes. LOTUS is a machine learning-based method that estimates a scoring function to rank candidate genes by decreasing probability that they are OGs or TSGs, given a training set of known OGs and TSGs. The score of a candidate gene is a weighted sum of similarities between the candidate gene and the known cancer genes, where the weights are optimized by a one-class support vector machine (OC-SVM) algorithm. The similarities themselves are derived from the analysis of somatic mutation patterns in the genes, or from the relative positions of genes in a PPI network, or from both; the mathematical framework of kernel methods allows to simply combine heterogeneous data about genes (i.e., patterns of somatic mutations and PPI information) in a single model.

Another salient feature of LOTUS is its ability to work in a pan-cancer setting, as well as to predict driver genes specific to individual cancer types. In the later case, we use a multitask learning strategy to jointly learn scoring functions for all cancer types by sharing information about known driver genes in different cancer types. We test both a default multitask learning strategy, that shares information in the same way across all cancer types, and a new strategy that shares more information across similar cancer types. More details about the mathematical formulation and algorithms implemented in LOTUS are provided in the Material and Methods section.

In the following, we assess the performance of LOTUS first in the pan-cancer regime, where we compare it to three state-of-the-art methods (TUSON, MutSigCV and 20/20+), and second in the cancer type specific regime, where we illustrate the importance of the multitask learning strategies.

### Cross-validation performance for pan-cancer driver gene prediction

We first study the pan-cancer regime where cancer is considered as a single disease, and where we search for driver genes involved in at least one type of cancer. Several computational methods have been proposed to solve this problem in the past, and we compare LOTUS with the three best methods in terms of performance according to a recent benchmark [27]: MutSigCV [21], which is a frequency-based method, and TUSON [25] and 20/20+ [27], which combine frequency and functional information.

While MutSigCV is and unsupervised method that scores candidate genes independently of any training set of known drivers, TUSON and 20/20+ depend on a training set, just like LOTUS. To perform a comparison as fair as possible between different methods, we collect the training sets of TUSON and 20/20+, and evaluate the performance of LOTUS on each of these datasets by 5-fold cross-validation (CV) repeated twice (see Methods). For TUSON and 20/20+, we use the prediction results available in the corresponding papers, in order to evaluate the consistency errors (*CE*) as the mean number of non-driver genes that are ranked before known driver genes of the TUSON and 20/20 train sets, respectively. We note that these ranks were obtained by training these two algorithms on their respective train set, and that this therefore gives an advantage to TUSON and 20/20+ compared to LOTUS in the evaluation.

Indeed for the former two methods the training set is used both to define the score and to assess the performance, while for LOTUS the CV procedure ensures that different genes are used to train the model and to test its performance. However we note that the 20/20+ score itself is obtained by a bootstrap procedure similar to our cross-validation approach [27]. This allows us to make fair comparisons between TUSON, MutSigCV and LOTUS (trained on TUSON train set), on the one hand, and between 20/20+, MutSigCV and LOTUS (trained on 20/20 train set), on the other hand. We further note that MutSigCV also provides a ranked list of genes, but does not make the difference between TSG and OG. Therefore, it is not dependent from a train set, and the *CE* in this case is obtained by averaging the numbers of non-driver genes ranked before each driver genes in the considered train set.

The *CE* for the different methods and the different training sets are presented in Table 1 for OGs and in Table 2 for TSGs. When analyzing these results, one should keep in mind that the total number of cancer driver genes is still a subject of debate, but it is expected to be much lower than the size of the test set of 17849 genes, and it should rather be in the range of a few hundreds. Therefore, consistency errors above a few thousand can be considered as poor performance results.

**Table 1.** Consistency error for OG prediction in the pan-cancer setting, for different methods (columns) and different gold standard sets of known OG (rows).

| Train set \ Method | MutSigCV | TUSON | 20/20+ | LOTUS |
| --- | --- | --- | --- | --- |
| TUSON train set | 4,489 | 3,286 | × | **931** |
| 20/20 train set | 5,823 | × | 1,831 | **819** |

**Table 2.** Consistency error for TSG prediction in the pan-cancer setting, for different methods (columns) and different gold standard sets of known TSG (rows).

| Train set \ Method | MutSigCV | TUSON | 20/20+ | LOTUS |
| --- | --- | --- | --- | --- |
| TUSON train set | 1,443 | 626 | × | **130** |
| 20/20 train set | 2,447 | × | 845 | **514** |

These results show that LOTUS strongly outperforms all other algorithms in term of *CE*, for both TSG and OG predictions. More precisly, for OG predictions, TUSON is about 5-fold better than MutSigCV, 3-fold better than TUSON and 2-fold better than 20/20+, in terms of *CE*. For TSG predictions, the reduction in *CE* with LOTUS is x, 5x and 1.6x compared to MutSigCV, TUSON and 20/20+, respectively. The performances of TUSON and 20/20+ are in the same range, although we should keep the above remark in mind. The results also show that MutSigCV does not perform as well as the three other methods, at least on the datasets used here.

It is interesting to note that, for all methods, the performances obtained for OG do not reach those obtained for TSG, suggesting that OG prediction is a more difficult problem than TSG prediction. This reflects the fundamental difference between TSG mutations and OG mutations: the first lead to loss-of-function and can pile up, while the second are gain-of-function mutations and have a much more subtle nature. In addition, gain-of-function can also be due to overexpression of the OG, which can arise from other mechanisms than gene mutation. One way to improve the OG prediction performance may be to include descriptors better suited to them, such as copy number. Moreover, as mutations affecting OGs are not all likely to provide them with new functionalities, many mutations on OGs present in the database and used here might not bear information on OGs. Therefore, relevant information on OGs is scarce, which makes OG prediction more difficult. In addition, the data themselves might also contribute to difference in performance between TSG and OG prediction. For example, in the case of the TUSON train set, although the TSG and OG train sets both contain 50 genes, the mutation matrix that we used to build the gene features contains 13, 525 mutations affecting TSGs and 7, 717 mutations affecting OGs. Therefore, the data are richer for TSG, which might contribute to the difference in prediction performance.

### The benefits of combining mutations and PPI informations

LOTUS, 20/20+, MutSigCV and TUSON differ not only by the algorithm they implement, but also by the type of data they use to make predictions: in particular, TUSON and 20/20+ use only mutational data while LOTUS uses PPI information in addition to mutational data. To highlight the contributions of the algorithm and of the PPI information to the performance of LOTUS, we ran LOTUS with *K*_*genes*_ = *K*_*mutation*_, or *K*_*genes*_ = *K*_*P* *P* *I*_, *i.e.*, with only mutation information, or only PPI information. The results are presented in Table 3 and Table 4 respectively for OG and TSG. The last column of these Tables recalls the performance obtained when mutation and PPI information are both used (values reported from Table 1 and Table 2).

**Table 3.** Consistency error of LOTUS for OG prediction in the pan-cancer setting, with different gene kernels (columns) and different gold standard sets of known OGs (rows).

| Train set \ Kernel | $K_{mutation}$ | $K_{PPI}$ | $K_{mutation} + K_{PPI}$ |
| --- | --- | --- | --- |
| TUSON train set | 2,333 | 1,565 | **931** |
| 20/20 train set | 2,072 | 2,013 | **819** |

**Table 4.** Consistency error of LOTUS for TSG prediction in the pan-cancer setting, with different gene kernels (columns) and different gold standard sets of known TSGs (rows).

| Train set \ Kernel | $K_{mutation}$ | $K_{PPI}$ | $K_{mutation} + K_{PPI}$ |
| --- | --- | --- | --- |
| TUSON train set | 388 | 1,645 | **130** |
| 20/20 train set | 901 | 1,858 | **514** |

These results show that, both for OG and TSG, using both mutation and PPI information dramatically improves the prediction performance over using only one type of them. This underlines the fact that mutation and PPI are complementary informations that are both useful for the prediction tasks. The performances obtained with only PPI information are similar for OG and TSG, which seems to indicate that this information contributes similarly to both prediction tasks. On the contrary, the performances obtained using only mutation information are much better for TSG than for OG. This is consistent with the above comment that mutation information is more abundant in the database and more relevant in nature for TSG than for OG. It is also consistent with the fact that using *K*_*mutation*_ alone outperforms using *K*_*P* *P* *I*_ alone for TSGs, while the opposite is observed for OGs.

### Performance on CGCv84 prediction in the pan-cancer regime

We now evaluate the generalization properties of the different methods on new unseen data as external test set. This not only mitigates the potential bias in the evaluation of the performance of TUSON and 20/20+ in the previous paragraph, but also allows to evaluate the performance of the different methods when predicting supposedly “difficult” new cancer genes, which have only been added recently in CGC. For that purpose we train LOTUS with the full 20/20 or TUSON train sets, make predictions on the full COSMIC database, and evaluate the *CE* using the CGCv84 database as a gold standard of true cancer genes, under the assumption that this database is enriched in driver genes (a criterion that was also used in [27]). We compare these *CE* to those of TUSON (for the TUSON train set) and 20/20+ (for the 20/20 train set). For LOTUS, TUSON and 20/20+, genes belonging to their corresponding trains set are removed from the CGCv84 database before calculating the *CE*. For MutSigCV, the *CE* is calculated based on the ranked list of genes provided in the corresponding paper [21], removing genes of the TUSON train set from CGCv84 database when MutSigCV is compared to TUSON and LOTUS (Table 5), and removing genes from the 20/20 train set from CGCv84 when MutSigCV is compared to 20/20+ and LOTUS (Table 6).

**Table 5.**
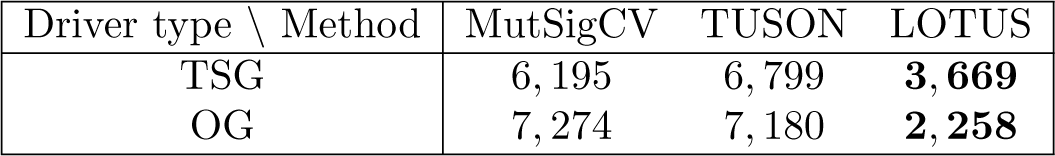
*CE* obtained on the CGCv84 data set with the TUSON train set.

**Table 6.** *CE* obtained on the CGCv84 data set with the 20/20 train set.

| Driver type \ Method | MutSigCV | 20/20+ | LOTUS |
| --- | --- | --- | --- |
| TSG | 6,925 | 4,893 | **3,944** |
| OG | 6,931 | 3,901 | **2,358** |

These results are illustrated by the corresponding ROC curves, see Figures 1 and 2.

**Fig 1.**
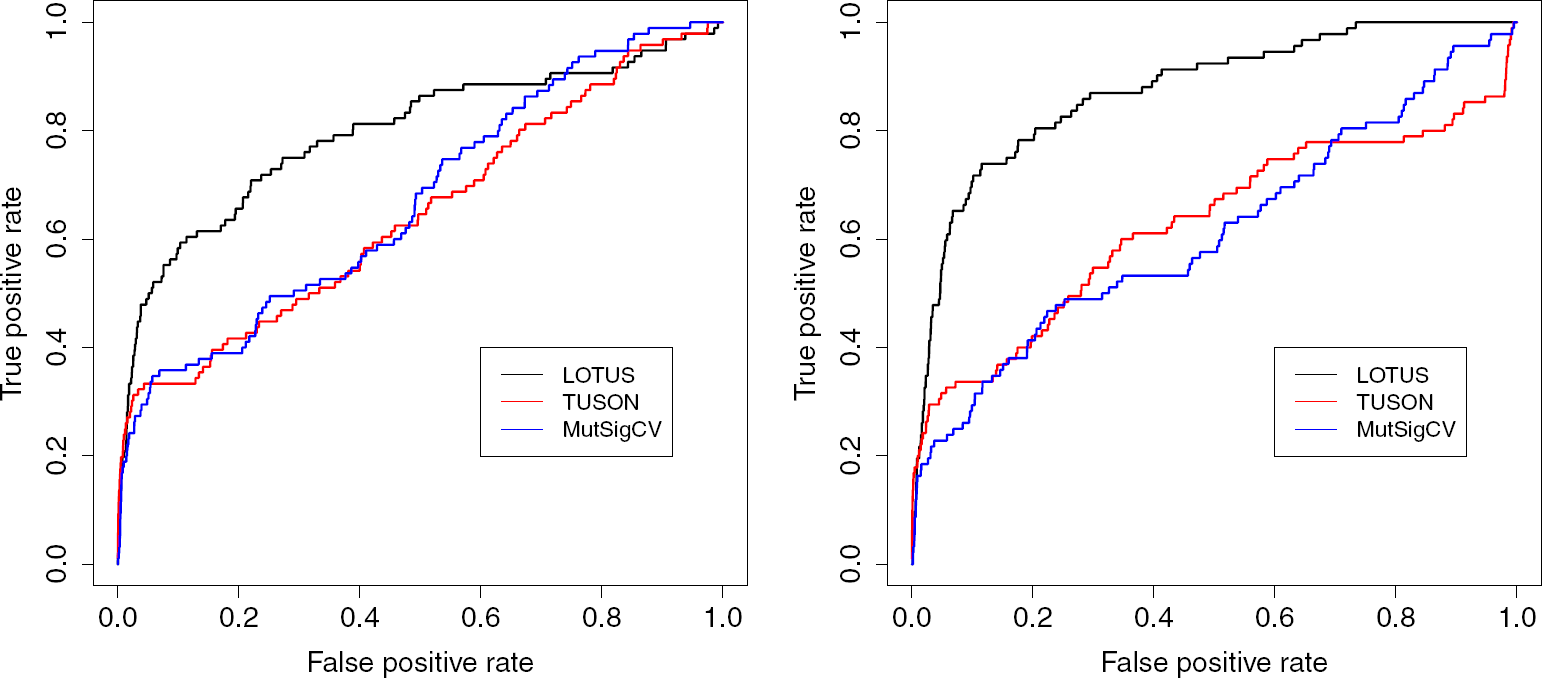
ROC curves for TSGs (left) and OGs (right) and the TUSON train set.

**Fig 2.**
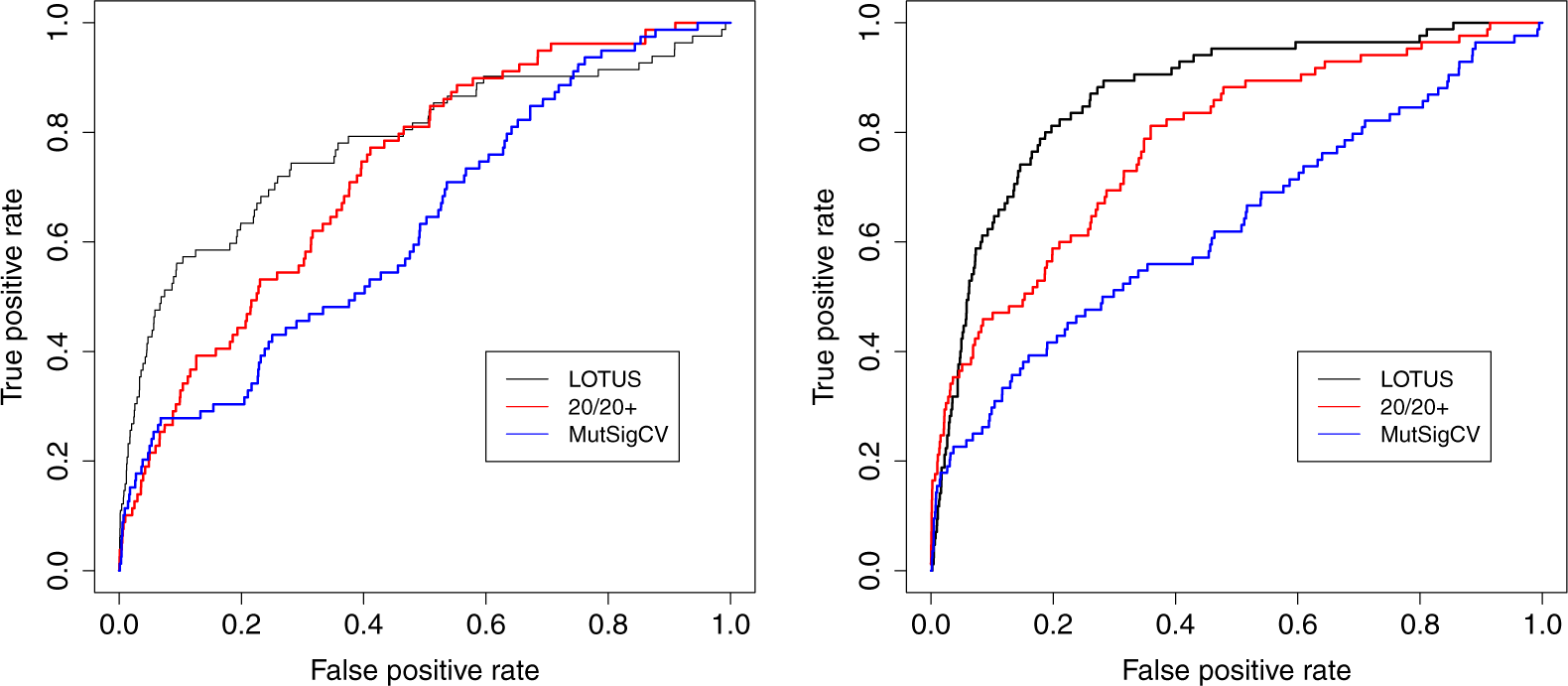
ROC curves for TSGs (left) and OGs (right) and the 20/20 train set.

We observe that, again, LOTUS strongly outperforms all three other methods in this setting. MutSigCV and TUSON have similar performance, and LOTUS outperforms them in all settings by a 1.6-to 3-fold decrease in *CE*. 20/20+ has better performance than MutSigCV, but has a *CE* 1.2 to 1.3 larger than LOTUS. We also observe that the absolute performance are overall worse than in the previous cross-validation experiment, which confirms the fact that genes recently added to CGC are overall harder to identify than the ones known for a long time.

### Analysis of new driver genes predicted by LOTUS

We now investigate the ability of LOTUS to make new driver gene predictions. For that purpose we train LOTUS with the CGCv84 train set, and make predictions over the complete COSMIC database (17, 948 genes). The complete results are given in Supplementary Table 3.

In the absence of experimental validation, we try to evaluate some of these predictions based on independent sources of information. Complete analyses of the predicted OG and TSG rankings is out of the scope of this paper. However, we consider below the 20 best ranked TSGs and OGs according to LOTUS.

Among the 20 best ranked TSGs, 4 genes are actually known TSGs that were not included yet in CGCv84: PTEN [32], FAT1 [33], STAG1 [34], TRAP1 [35].

Interestingly, 8 genes out of these 20 best ranked TSGs are genes coding for proteins involved in DNA repair, a role closely related to genome maintenance and cancer [36, 37]. These genes are EXO1 [38], ERCC1 [39], GTF2H1 and GTF2H4 (both involved in the TFIIH complex [40]), NTHL1 [41], ATR [42], RAD52 [43] and RPA4 [44]. In addition to these clues referring to the DNA repair functions, many additional studies related to these genes are available in the literature, underlining their role in various types of cancers, which provides another clue for them to be confident TSG candidates. In particular, mutations in NTHL1 are known to predispose to colorectal cancer, which is an additional argument in favor of NTHL1 being a strong candidate TSG [45, 46].

For 2 additional genes, GALNT5 and PIWIL1, we find recent publications indicating that they could potentially act as TSG, at least in some tumor types. A non-coding RNA directed against GALNT5 is overexpressed in gastric cancer, inhibiting the translation of its target gene, and the level of expression of this non-coding RNA is correlated with cancer progression and metastasis [47]. These results are consistent with a TSG role of GALNT5 in gastric cancer. In the case of PIWIL1, a recent paper concludes that it is an epidriver gene for lung adenocarcinoma, which means that aberrant methylation of its promoter region plays a role in the development of this cancer [48].

Among the 20 best ranked putative OGs, 3 genes are actually known OGs at least for some types of cancers, and not yet included in CGCv84: MAP3K1 [49], PLCE1 [50], FGF5 [51].

One gene, GATA3, is known to behave either as an OG or as a TSG, depending on the genetic context of the disease [52]. In fact, the literature provides other examples of genes able to switch from oncogenes to tumor suppressor genes, depending on the context [53]. In line with this remark, 3 genes among the 20 best ranked OGs are known TSGs. They could in fact have a potential property to be OG or TSG, depending on the context: PIK3R1 [54], APC [55], TP53 [56].

Mutations in the 6th ranked HTPO gene seems to be causal in some cancer types, where it could therefore be considered as an oncogene [57].

Finally 4 genes are known to be associated to cancer development and progression in some cancer types, are studied as biomarkers or as therapeutic targets, which indicates that they could indeed be credible oncogene candidates: PPARP10 [58], HTR2B [59], STAP2 [60], FXYD2 [61].

Taken together, these results show that LOTUS is able to retrieve, among the top ranked genes, known driver genes that are absent from the training set. They also show that LOTUS suggests high confidence driver genes for which many references about their implication in cancer are available.

### Identification of cancer-specific driver genes with multitask LOTUS

In this section, we do not consider cancer as a single disease, but as a variety of diseases with different histological types and that can affect various organs. It is then important to identify driver genes for each type of cancer. One way to solve this problem is to use a prediction method that is trained only with driver genes known for the considered cancer. Such single-task methods may however display poor performance because the number of known drivers per cancer is often too small to derive a reliable model. Indeed, scarce training data lead to a potential loss of statistical power as compared to the problem of identification of pan-cancer driver genes were data available for all cancers are used.

In this context, we investigate the multitask versions of LOTUS, where we predict driver genes for a given cancer based on the drivers known for this cancer but also on all driver genes known for other cancer types. For a given cancer type, this may improve driver genes prediction by limiting the loss of statistical power compared to the aforementioned single-task approach.

For that purpose, we derive a list of 174 cancer diseases from COSMICv84 as explained in Methods. This complete list is available in Supplementary Table 1. As expected, many cancer types have only few, if any, known cancer genes (Figure 3).

**Fig 3.**
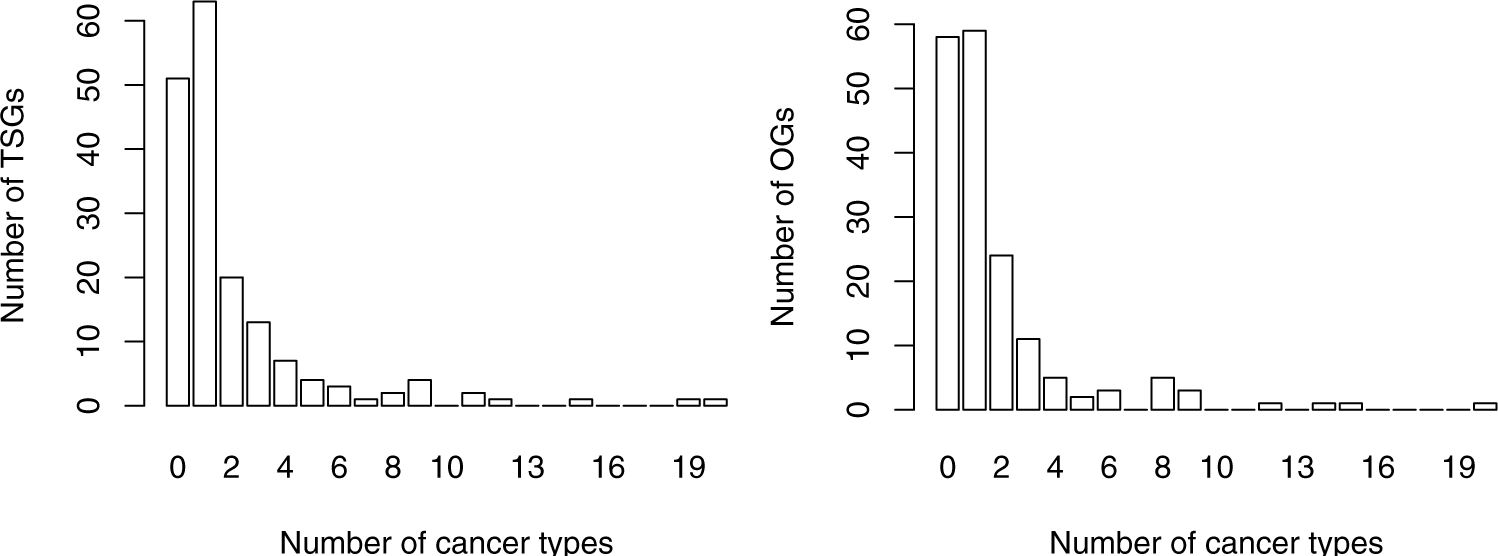
Distribution of the number of TSGs (left) and OGs (right) per cancer type.

Since we want to evaluate the performance of LOTUS in a cross-validation scheme, we only consider diseases with more than 4 known driver genes in order to be able to run a 2-fold CV scheme. This leads us to keep 27 cancer types for TSG prediction and 22 for OG prediction. Note however that prediction are made for these 27 and 22 cancer types while sharing all the driver genes known for the 174 diseases (according to their similarities with these 27 and 22 cancer types).

The 2-fold CV consistency error of LOTUS for each of those cancer types is presented in Tables 7 (for TSG) and 8 (for OG). Here we compare four variants of LOTUS, as explained in Methods: single-task LOTUS treats each disease in turn independently from the others; aggregation LOTUS applies a pan-cancer prediction by pooling together the known genes of all cancer types; and the two multitask versions of LOTUS use either a standard multitask strategy that do not take into account the relative similarities between diseases (multitask TUSON), or a more refined multitask strategy where similar cancer types share more information than non-similar ones (multitask TUSON2).

**Table 7.**
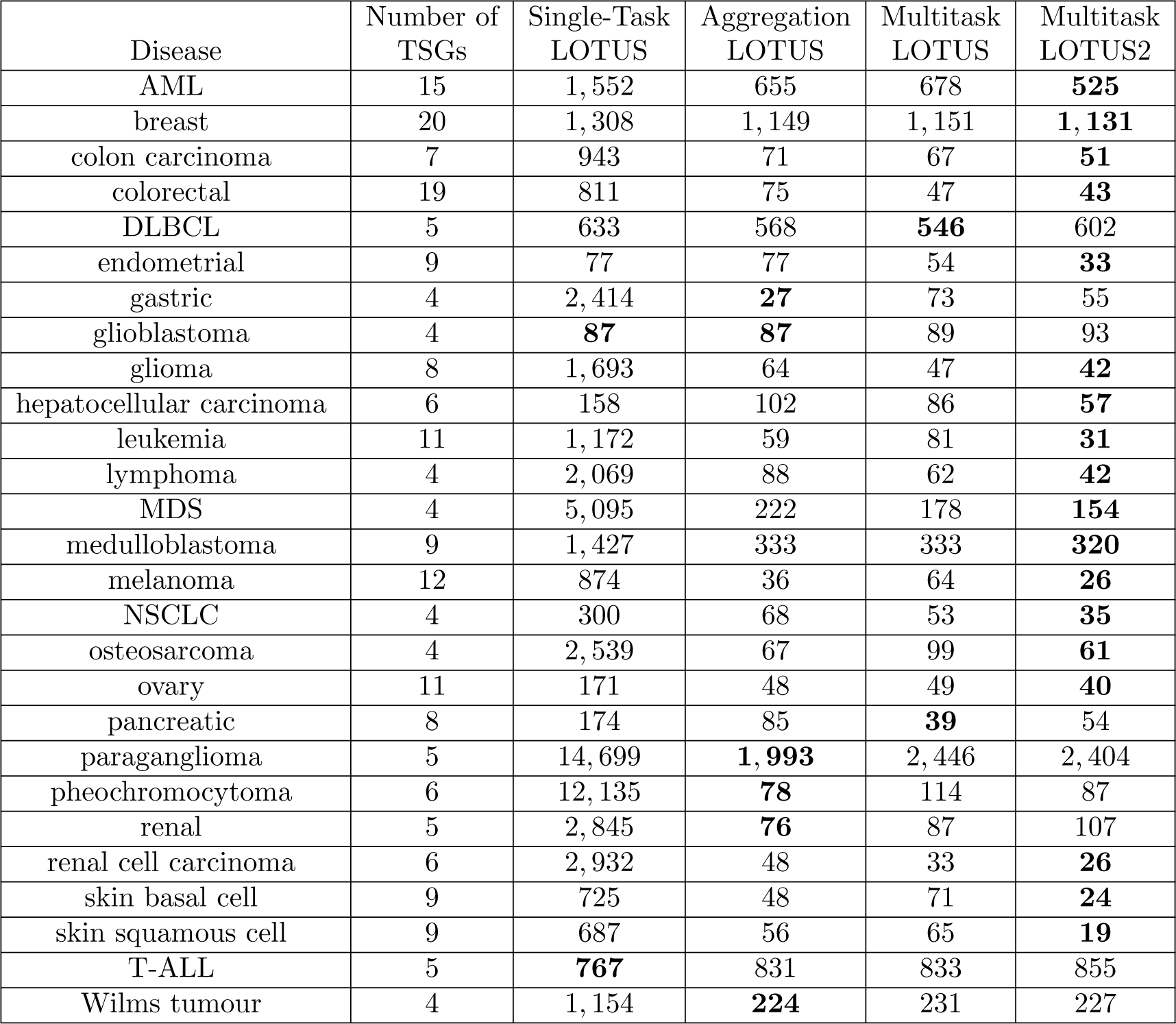
*CE* for prediction of disease specific TSGs in the multitask setting.

| Disease | Number of TSGs | Single-Task LOTUS | Aggregation LOTUS | Multitask LOTUS | Multitask LOTUS2 |
| --- | --- | --- | --- | --- | --- |
| AML | 15 | 1,552 | 655 | 678 | **525** |
| breast | 20 | 1,308 | 1,149 | 1,151 | **1,131** |
| colon carcinoma | 7 | 943 | 71 | 67 | **51** |
| colorectal | 19 | 811 | 75 | 47 | **43** |
| DLBCL | 5 | 633 | 568 | **546** | 602 |
| endometrial | 9 | 77 | 77 | 54 | **33** |
| gastric | 4 | 2,414 | **27** | 73 | 55 |
| glioblastoma | 4 | **87** | **87** | 89 | 93 |
| glioma | 8 | 1,693 | 64 | 47 | **42** |
| hepatocellular carcinoma | 6 | 158 | 102 | 86 | **57** |
| leukemia | 11 | 1,172 | 59 | 81 | **31** |
| lymphoma | 4 | 2,069 | 88 | 62 | **42** |
| MDS | 4 | 5,095 | 222 | 178 | **154** |
| medulloblastoma | 9 | 1,427 | 333 | 333 | **320** |
| melanoma | 12 | 874 | 36 | 64 | **26** |
| NSCLC | 4 | 300 | 68 | 53 | **35** |
| osteosarcoma | 4 | 2,539 | 67 | 99 | **61** |
| ovary | 11 | 171 | 48 | 49 | **40** |
| pancreatic | 8 | 174 | 85 | **39** | 54 |
| paraganglioma | 5 | 14,699 | **1,993** | 2,446 | 2,404 |
| pheochromocytoma | 6 | 12,135 | **78** | 114 | 87 |
| renal | 5 | 2,845 | **76** | 87 | 107 |
| renal cell carcinoma | 6 | 2,932 | 48 | 33 | **26** |
| skin basal cell | 9 | 725 | 48 | 71 | **24** |
| skin squamous cell | 9 | 687 | 56 | 65 | **19** |
| T-ALL | 5 | **767** | 831 | 833 | 855 |
| Wilms tumour | 4 | 1,154 | **224** | 231 | 227 |

**Table 8.** *CE* for prediction of disease specific OGs in the multitask setting

| Disease | Number of<br>OGs | Single-Task<br>LOTUS | Aggregation<br>LOTUS | Multitask<br>LOTUS | Multitask<br>LOTUS2 |
| --- | --- | --- | --- | --- | --- |
| ALL | 9 | 1,637 | 873 | 856 | **796** |
| AML | 20 | 1,447 | 606 | 600 | **578** |
| bladder | 5 | 636 | 83 | **32** | 54 |
| breast | 8 | 2,250 | 121 | 134 | **91** |
| CLL | 8 | 2,598 | 824 | **814** | 825 |
| colorectal | 12 | 2,018 | 68 | 32 | **27** |
| DLBCL | 5 | **107** | 355 | 353 | 327 |
| endometrial | 6 | 616 | 40 | 28 | **26** |
| gastric | 9 | 112 | 40 | 25 | **15** |
| glioblastoma | 8 | 3,452 | 74 | 60 | **54** |
| glioma | 6 | **613** | 761 | 773 | 769 |
| head and neck | 6 | 320 | 71 | 51 | **39** |
| lymphoma | 4 | 5,651 | 79 | **61** | 77 |
| MDS | 9 | 5,071 | 86 | 109 | **82** |
| melanoma | 14 | 1,420 | 281 | **276** | 295 |
| MM | 4 | 3,122 | 77 | **37** | 60 |
| NSCLC | 15 | 2,281 | 280 | **126** | 149 |
| ovary | 8 | 3,194 | 57 | 37 | **32** |
| prostate | 8 | 845 | 162 | **126** | 154 |
| Spitzoid tumour | 4 | 183 | 68 | **38** | 48 |
| T-ALL | 4 | 8,436 | **2,041** | 2,047 | 2,046 |
| WM | 4 | 203 | 162 | 160 | **78** |

For most diseases (25/27 for TSG, 20/22 for OG), single-task LOTUS leads to the worst *CE*, confirming the difficulty to treat each cancer type individually due to the small number of know cancer gene for each individual type. Interestingly, Aggregation LOTUS often leads to a strong improvement in performance. This shows that different cancer types often share some common mechanisms and driver genes, and therefore, simply using all the available information in a pan-cancer paradigm improves the performance of driver gene prediction for each cancer type. However, in many cases, the multitask LOTUS and LOTUS2 algorithms lead to an additional improvement over Aggregation LOTUS, LOTUS2 leading in general to the best results (in 18 types out of 27 for TSG prediction, and in 11 types out of 22 for OG prediction). On average, the decrease in *CE* between Aggregate LOTUS and LOTUS2 is of 23% for OG and 17% for TSG. The improvement in performance observed between Aggregate LOTUS and LOTUS2 shows that, besides some driver mechanisms common to many cancers, each cancer presents some specific driver mechanisms that can only be captured by prediction methods able to integrate some biological knowledge about the diseases. The above results show that multitask algorithms allowing to share information between cancers according to their biological similarities such as LOTUS2, rather than on more naive rules, better capture these specific driver genes. They also show that the kernel *K*_*diseases*_ = *K*_*descriptors*_ built on disease descriptors contains some relevant information to compare diseases.

In the above table, AML stands for acute myeloid leukemia, DLBCL for diffuse large B-cell lymphoma, MDS for myelodysplastic syndromes, NSCLC for non-small cell lung cancer and T-ALL for T-cell acute lymphoblastic cancer.

Taken together, these results show that multitask machine learning algorithms like LOTUS are interesting approaches to predict cancer specific driver genes. In addition, multitask algorithms based on task descriptors (here, disease descriptors) appear to be promising in order to include prior knowledge about diseases and share information according to biological features characterizing the diseases.

In the above table, ALL stands for acute lymphotic leukemia, AML for acute myeloid leukemia, CLL for chronic lymphocytic leukemia, DLBCL for diffuse large B-cell lymphoma, MDS for myelodysplastic syndromes, MM for multiple myeloma, NSCLC for non-small cell lung cancer, T-ALL for T-cell acute lymphoblastic cancer and WM for Waldenstrom macroglogulinemia.

Finally, note that we did not try to run TUSON, MutSigCV or 20/20+ to search for cancer specific driver genes. Indeed, according to the results of pan-cancer studies in the single-task setting, they do not perform as well as single-task LOTUS. Moreover, they are not adapted, as such, to the multitask setting.

## Discussion

Our work demonstrates that LOTUS outperforms several state-of-the-art methods on all tested situations for driver gene prediction. This improvement results from various aspects of the LOTUS algorithm. First, LOTUS allows to include the PPI network information as independent prior biological knowledge. In the single-task setting, we proved that this information has significance for the prediction of cancer driver genes. Because LOTUS is based on kernel methods, it is well suited to integrate other data from multiple sources such as protein expression data, information from chip-seq, HiC or methylation data, or new features for mutation timing as designed in [62]. Further development could involve the definition of other gene kernels based on such type of data, and combine them with our current gene kernel, in order to evaluate their relevance in driver gene prediction.

We also showed how LOTUS can serve as a multitask method. It relies on a disease kernel that controls how driver gene information is shared between diseases. Interestingly, we showed that building a kernel based on independent biological prior knowledge about disease similarity leads on average to the best prediction performance with respect to single-task algorithms, and also with respect to a more generic multitask learning strategy that does not incorporate knowledge about the cancer types. Again, the kernel approach leaves space for integration of other types and possibly more complex biological sources of information about diseases. Our multitask approach thus allows to make prediction for cancer types with very few known driver genes, which would be less reliable with the single-task methods. We considered here only diseases with at least 4 known driver genes, in order to perform cross-validation studies, which was necessary to evaluate the methods. However, it is important to note that in real-case studies, at the extreme, both versions of multitask LOTUS could make driver gene prediction for cancer types for which no driver gene is known.

Among the 174 diseases derived form the COSMIC database, we kept only 27 cancer types for TSG prediction and 22 for OG prediction, for which at least four driver genes were available. However, inspection of the 174 disease names indicates that there might be diseases that could be grouped (for example “colorectal” and “colorectal adenocarcinoma”, or “skin” with “skin basal cell” or “skin squamous cell”), which would have allowed to enlarge the training sets and possibly improve the predictions. Future directions could be to have experts analyze and potentially modify this disease list, in order to optimize the training sets, or help to derive finer disease descriptors.

LOTUS is a machine learning algorithm based on one-class SVM. In fact, the most classical problem in machine learning is binary classification, where the task is to classify observations into two classes (positives and negatives), based on training sets *𝒫* of known positives and *𝒩* of known negatives. Driver gene detection can be seen as binary classification of TSGs vs. neutral genes, and of OGs vs. neutral genes. However, although the *𝒫* set is composed of known driver genes, it is not straightforward to build the *𝒩* set because we cannot claim that some genes cannot be drivers. Thus, driver gene detection should rather be seen as binary classification problem with only one training set *𝒫* of known positives. This problem is called classically called PU learning (for Positive-Unknown), as opposed to PN learning (for Positive-Negative).

The classical way to solve PU learning problems is to choose a set *𝒩* of negatives among the unlabeled data and apply a PN learning method. For example, one can consider all unknown items as negatives (some of which may be reclassified afterwards as positives), or randomly choose bootstrapped sets of negatives among the unknown, like in [31]. Both methods assume that a minority of the unlabeled items are in fact positives, which is expected for driver genes.

The one-class SVM algorithm [63] can also be used as a PU learning method, in which a virtual item is chosen as the training set of negatives. We preferred this approach because in preliminary studies, we found that it had slightly better performances than PU learning methods and was also faster.

For LOTUS, as for all machine learning algorithm, the set of known driver genes is critical: if this set is poorly chosen (*i.e.*, if some genes were wrongly reported as driver genes, or more likely, if the reported genes are not the best driver genes), the best algorithm might not minimize the consistency error *CE*. To circumvent this problem, we propose two new approaches for future developments: one could build a multi-step algorithm that iteratively removes some genes from the positive set and labels them as unknown, and add relabel as positives some of the best ranked unknown genes. We believe that such an algorithm would make the set of positives converge to a more relevant list. Alternatively, one could assign (finite) scores to the known driver genes before performing classification and increment these scores at each step.

## Materials and methods

### Pan-cancer LOTUS

LOTUS is a new machine learning-based method to predict new cancer genes, given a list of know ones. In the simplest, pan-cancer setting, we thus assume given a list of *N* known cancer genes {*g*_1_, *…, g_N_*}, and the goal of LOTUS is to learn from them a scoring function *f* (*g*), for any other gene *g*, that predicts how likely it is that *g* is also a cancer gene. Since TSGs and OGs have different characteristics, we treat them separately and build in fact two scoring functions *f*_*T* *SG*_ and *f*_*OG*_ trained from lists of know TSGs and OGs, respectively.

LOTUS learns the scoring function *f* (*g*) with a one-class support vector machine (OC-SVM) algorithm [63], a classical method for novelty detection and density level set estimation [64]. The scoring function *f* (*g*) learned by a OC-SVM given a training set {*g*_1_, *…, g_N_*} of known cancer genes takes the form:

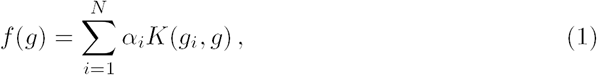

where *a*_1_, *…, a_N_* are weights optimized during the training of OC-SVM [63], and *K*(*g, g^′^*) is a so-called *kernel* function that quantifies the similarity between any two genes *g* and *g*^*′*^. In other words, the score of a new gene *g* is a weighted combination of its similarities with the know cancer genes.

The kernel *K* encodes the similarity among genes. Mathematically, the only constraint that *K* must fulfill is that it should be a symmetric positive definite function [29]. This leaves a lot of freedom to create specific kernels encoding one’s prior knowledge about relevant information to predict cancer genes. In addition, one can easily combine heterogeneous information in a single kernel by, e.g., summing together two kernels based on different sources of data. In this work, we restrict ourselves to the following basic kernels, and leave for future work a more exhaustive search of optimization of kernels for cancer gene prediction.

- *Mutation kernel.* Given a large data set of somatic mutations in cohorts of cancer patients, we characterize each gene *g* by a vector Φ_*mutation*_(*g*) ∈ ℝ_3_ encoding 3 features. For OG prediction the three features are the number of damaging missense mutations, the total number of missense mutations, and the entropy of the spatial distribution of the missense mutations on each gene. For TSG prediction, the features are the number of frameshift mutations, the number of LOF mutations (defined as the nonsense and frameshift mutations), and the number of splice site mutations. These features were calculated as proposed by [25]. We chose them because they were found to best discriminate OGs and TSGs by the TUSON algorithm [25] and were also all found among the most important features selected by the random forest algorithm used by the 20/20+ method [27]. Given two genes *g* and *g*^*′*^ represented by their 3-dimensional vectors Φ(*g*) and Φ(*g*^*′*^), we then define the mutation kernel as the inner product between these vectors:

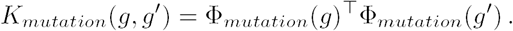

Notice that using *K*_*mutation*_ as a kernel in OC-SVM produces a scoring function (1) which is simply a linear combination of the three features used to define the vector Φ_*mutation*_.
- *PPI kernel.* Given an undirected graph with genes as vertices, such as a PPI network, we define a PPI kernel *K*_*P P I*_ as a graph kernel over the network [65, 66]. More precisely, we used a diffusion kernel of the form *K*_*P P I*_ = exp*M* (*-L*), where *L* = *I* - *D*^−1/2^*AD*^−1/2^ is the normalized Laplacian of the graph and exp*M* is the matrix exponential function. Here *I* is the identity matrix, *A* stands for the adjacency matrix of the graph (*A*_*i,j*_ = 1 if vertices *i* and *j* are connected, 0 otherwise) and *D* for the diagonal matrix of degrees 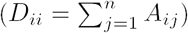. Intuitively, two genes are similar according to *K*_*P P I*_ when they are close and well connected through several routes to each other on the PPI network, hence learning a OC-SVM with *K*_*P P I*_ allows to diffuse the information about cancer genes over the network.
- *Integrated kernel.* In order to train a model that incorporates informations about both mutational features and PPI, we create an integrated gene kernel by simply averaging the mutation and PPI kernels:

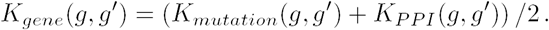

While more complex kernel combination strategies such as multiple kernel learning could be considered, we restrict ourselves to this simple kernel addition scheme to illustrate the potential of our approach for heterogeneous data integration.

### Multitask LOTUS for cancer type-specific predictions

The pan-cancer LOTUS approach can also be used for cancer-specific predictions, by restricting the training set of known cancer genes to those cancer genes known to be driver in a particular cancer type. However, for many cancer types, only few driver genes have been validated, creating a challenging situation for machine learning-based methods like LOTUS that rely on a training set of known genes to learn a scoring function. Since cancer genes of different cancer types are likely to have similar features, we propose instead to learn jointly cancer type-specific scoring functions by sharing information about known cancer genes across cancer types, using the framework of multitask learning [30, 31]. Instead of starting from a list of known cancer genes, we now start from a list of known (cancer gene, cancer type) pairs of the form {(*g*_1_, *d*_1_), *…,* (*g*_*N*_, *d*_*N*_)}, where a sample (*g*_*i*_, *d*_*i*_) means that gene *g*_*i*_ is a known cancer gene in disease *d*_*i*_. Note that a given gene (and a given cancer type) may of course appear in several such pairs.

The extension of OC-SVM to the multitask setting is straightforwardly obtained by creating a kernel for (gene, disease) pairs of the form:

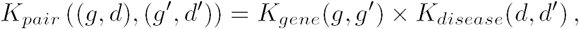

where *K*_*gene*_ is a kernel between genes such as the one used in pan-cancer LOTUS and *K*_*disease*_ is a kernel between cancer types described below. We then simply run the OC-SVM algorithm using *K*_*pair*_ as kernel and {(*g*_1_, *d*_1_), *…,* (*g*_*N*_, *d*_*N*_)} as training set, in order to learn a cancer type-specific scoring function of the form *f* (*g, d*) that estimates the probability that *g* is a cancer gene for cancer type *d*.

The choice of the disease kernel *K*_*disease*_ influences how information is shared across cancer types. One extreme situation is to take the uniform kernel *K*_*uniform*_(*d, d^′^*) = 1 for any *d, d^′^*. In that case, no distinction is made between diseases, and all known cancer genes are pooled together, recovering the pan-cancer setting (with the slight difference that genes may be counted several times in the training set if they appear in several diseases). Another extreme situation is to take the Dirac kernel *K*_*Dirac*_(*d, d^′^*) = 1 if *d* = *d*^*′*^, 0 otherwise. In that case, no information is shared across cancer types, and the joint model over (gene, disease) pairs is equivalent to learning independently a model for each disease.

In order to leverage the benefits of multitask learning and learn disease-specific models by sharing information across diseases, we consider instead the following two disease kernels:

- First, we consider the standard multitask learning kernel:

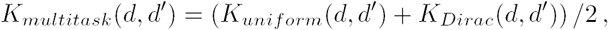

which makes a compromise between the two extreme uniform and Dirac kernels [30]. Intuitively, for a given cancer type, prediction of driver genes is made by assigning twice more weight to the data available for this cancer than to the data available for all other cancer types.
- Second, we test a more elaborate multitask version where we implement the idea that a given cancer might share various degrees of similarities with other cancers. Therefore, known cancer genes for other cancers should be shared with those of the considered cancer based on this similarity. Hence we create a specific disease kernel *K*_*cancer*_(*d, d^′^*) to capture our prior hypothesis about how similar cancer genes are likely to be between different cancers. To create *K*_*cancer*_, we first represent each cancer type as a 50-dimensional binary vector as follows. The first 15 bits correspond to a list of cancer type characteristics used in COSMIC to describe tumors: adenocarcinoma, benign, blastoma, carcinoma, gastro-intestinal stromal tumour, germ cell tumour, glioma, leukemia, lymphoma, melanoma, meningioma, myeloma, neuro-endocrine, sarcoma, stromal. The last 35 components correspond to localization characteristics also used in COSMIC to describe tumors: bile ducts, bladder, blood vessels, bone, bone marrow, breast, central nervous system, cervix, colorectal, endocrine glands, endometrium, eye, gall bledder, germ cell, head and neck, heart, intestine, kidney, liver, lung, lymphocytes, mouth, muscle, nerve, oesophagus, ovary, pancreas, pituitary glands, prostate, salivary glands, skin, soft tissue, stomach, tendon, thyroid. A disease might be assigned one or several types and be associated to one or several locations. For example, neurofibroma is associated with a single localization (“nerve”) and two types (“benign” and “sarcoma”), so that neurofibroma is described by a vector with three 1’s and forty-seven 0’s. For each disease, we construct the list of binary features by documenting every disease in the literature. The corresponding vectors encoding the considered disease are given in Supplementary Table S2. Finally, if Ψ(*d*) *∈* ℝ_50_ denotes the binary vector representation of disease *d*, we create the disease kernel as a simple inner product between these vectors, combined with the standard multitask kernel, i.e.:

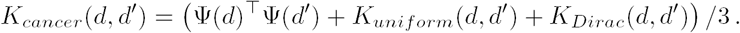

### Data

In all experiments, we restrict ourselves to the total set of 17, 948 genes considered in the TUSON, 20/20 and MutSigCV papers, as candidate driver genes. Somatic mutations were collected from COSMIC [14], TCGA (http://cancergenome.nih.gov/) and [18]. This dataset contains a total of 1, 195, 223 mutations across 8, 207 patients. We obtained the PPI network from the HPRD database release 9 from April 13, 2010 [67]. It contains 39, 239 interactions among 7, 931 proteins. As for known pan-cancer driver genes, we consider three lists in our experiments: (i) the TUSON train set, proposed in [25], consists of two high confidence lists of 50 OGs and 50 TSGs extracted from CGC (release v71) based on several criteria, in particular excluding driver genes reported through translocations; (ii) the 20/20 train set, proposed in [27] to train the 20/20+ method, contains 53 OGs and 60 TSGs; finally, (iii) the CGCv84 train set consists of two broader lists that we extracted from CGC release v84 of the COSMIC database: the list of all 136 dominant driver genes in the CGC database that were not reported through translocations (i.e., OGs), and the list of all 138 recessive driver genes in the CGC database that were not reported through translocations (i.e., TSGs). For cancer type-specific lists of driver genes, we only consider the CGCv84 train set. We distinguished 174 diseases based on the available annotations describing patients in COSMIC, using as few interpretations as possible: for example, we merged together diseases corresponding to obvious synonyms like singular and plural forms of the same cancer name. The names of these diseases and their numbers of associated TSGs and OGs can be found in Supplementary Table 1. For each of the resulting diseases, 1 to 20 TSGs/OGs were known in CGCv84. We considered only diseases with at least 4 known TSGs or OGs available, in order to have enough learning data points to perform a cross-validation scheme, which led us to consider 27 diseases for TSG prediction and 22 for OG prediction.

### Experimental protocol

To assess the performance of a driver gene prediction method on a given gold standard of known driver genes, we score all genes in the COSMIC database and measure how well the known driver genes are ranked. For that purpose, we plot the receiver operating characteristic (ROC) curve, considering all known drivers as positive examples and all other genes in COSMIC as negative ones, and define the consistency error (*CE*) as

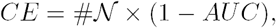

where #*N* is the number of negative genes, and *AUC* denotes the area under the ROC curve. In words, *CE* measures the mean number of “non-driver” genes that the prediction method ranks higher than known driver genes. Hence, a perfect prediction method should have *CE* = 0, while a random predictor should have a *CE* near #*N /*2.

To estimate the performance of a machine learning-based prediction method that estimates a scoring function from a training set of known driver genes, we use *k*-fold cross-validation (CV) for each given gold standard set of known driver genes. In *k*-fold CV, the gold standard set is randomly split into *k* subsets of roughly equal sizes. Each subset is removed from the gold standard in turn, the prediction method is trained on the remaining *k -*1 subsets, and its *CE* is estimated considering the subset left apart as positive examples, and all other genes of COSMIC not in the gold standard set as negative examples. A mean ROC curve and mean *CE* is then computed from the *k* resulting ROC curves. This computation is repeated several times to consider several possibly different partitions of the gold standard set.

### Tuning of parameters

Each version of LOTUS depends on a unique parameter, the regularization parameter *C* of the OC-SVM algorithm. Each time a LOTUS model is trained, its *C* parameter is optimized by 5-fold CV on the training set, by picking the value in a grid of candidate values {2^−5/2^, 2^−4/2^, …, 2^5/2^} that minimizes the mean *CE* over the folds.

### Other driver prediction methods

We compare the performance of LOTUS to three other state-of-the-art methods: MutSigCV [21], which is a frequency-based method, and TUSON [25] and 20/20+ [27] that combine frequency and functional information.

MutSigCV searches driver genes among significantly mutated genes which adjusts for known covariates of mutation rates. The method estimates a background mutation rate for each gene and patient, based on the observed silent mutations in the gene and noncoding mutations in the surrounding regions. Incorporating mutational heterogeneity, MutSigCV eliminates implausible driver genes that are often predicted by simpler frequency-based models. For each gene, the mutational signal from the observed non-silent counts are compared to the mutational background. The output of the method is an ordered list of all considered genes as a function of a p-value that estimates how likely this gene is to be a driver gene.

TUSON uses gene features that encode frequency mutations and functional impact mutations. The underlying idea is that the proportion of mutation types observed in a given gene can be used to predict the likelihood of this gene to be a cancer driver. After having identified the most predicting parameters for OGs and TSGs based on a train set (called the TUSON train set in the present paper), TUSON uses a statistical model in which a p-value is derived for each gene that characterizes its potential as being an OG or a TSG, then scores all genes in the COSMIC database, to obtain two ranked lists of genes in increasing orders of p-values for OGs and TSGs.

The 20/20+ method encodes genes based on frequency and mutation types, and other biological information. It uses a train set of OGs and TSGs (called the 20/20 train set in the present paper) to train a random forest algorithm. Then, the random forest is used on the COSMIC database and the output of the method is again a list of genes ranked according to their predicted score to be a driver gene [27]. We did not implement this method, so we decided to evaluate its performance only on its original training set: the 20/20 dataset. Moreover, we applied the same method to compute the *CE* as for MutSigCV and TUSON, which should actually give an advantage to 20/20+, since it is harder to make predictions in a cross-validation loop using a smaller set of known driver genes.

### Code and data availability

We implemented LOTUS and performed all experiments in R using in particular the kernlab package for OC-SVM [68]. The code and data to reproduce all experiments are available at http://members.cbio.mines-paristech.fr/∼ocollier/lotus.html.

## Acknowledgments

This work was supported the European Research Council grant ERC-SMAC-280032 (OC, JPV) and the Labex MME-DII ANR11-LBX-0023-01 (OC).

## Supporting information

**S1 Table List of cancer types (CGC v84).** Cancer types derived from COSMIC annotations along with their numbers of associated OG and TSG. The resulting names are sometimes very general and sometimes very specific, and some redundancies may be present, because we chose to add as little interpretation as possible.

**S2 Table Description of cancer types (CGC v84).** Descriptors of all cancer types according to their localizations and types that are used to compute the disease kernel used by LOTUS2.

**S3 Table TSG and OG rankings for LOTUS with the 20/20, the TUSON and the CGCv84 datasets.** Note that the training sets were removed every time.

